# An integrated virtual reality platform for naturalistic neuroimaging with magnetoencephalography

**DOI:** 10.64898/2026.01.28.701672

**Authors:** Sajjad Zabbah, Nicholas A Alexander, Yousef Mohammadi, Alberto Mariola, Robert A Seymour, Sahitya Puvvada, Gareth R Barnes, Dominik R Bach

## Abstract

Studying the brain in motion promises deep insights into the neural circuits that support complex, real-world behaviour. In humans, wearable optically pumped magnetometers (OPMs) enable magnetoencephalography (MEG) with millisecond temporal resolution and millimetre spatial precision during movement. Integrating this technology with virtual reality (VR) could enable fully naturalistic experimental paradigms, but magnetic interference from existing head-mounted displays (HMDs) prevents reliable whole-brain MEG recordings. Here, we present and validate a VR system that integrates with wearable, OPM-based MEG. At its core is a purpose-designed HMD with minimal ferromagnetic material, resulting in magnetic flux density two orders of magnitude lower than consumer-grade alternatives at comparable resolution and weight. Using phantom measurements and established perceptual and cognitive benchmark tasks across participants, we demonstrate robust stimulus-induced neuronal activity at both sensor and source level. Crucially, these sources span a broad set of brain regions, including visual, motor and prefrontal cortices. There remain significant noise peaks which will challenge certain experimental designs however, the system is straightforward to produce, readily extendable, and enables whole-brain MEG during immersive, naturalistic behaviours.

**Significance statement:** Understanding the human brain during natural behaviour requires recording neural activity while people move and interact with realistic environments. Virtual reality offers a powerful means to simulate such environments in a controlled setting, but existing headsets generate magnetic interference that prevents their use with wearable brain scanners based on optically pumped magnetometers, the only neuroimaging technology capable of whole-brain recording during free movement. We designed and validated a low-interference headset that reduces magnetic noise by two orders of magnitude compared to commercial alternatives, enabling whole-brain recordings spanning visual, motor and prefrontal regions. This open-source platform is straightforward to build and extendable, establishing a foundation for studying human cognition in controllable, realistic environments with millisecond and millimetre precision.

## Introduction

Non-invasive human neuroimaging has enabled fundamental insights into the neural mechanisms underlying perception, action, and cognition. Yet, most established neuroimaging modalities impose severe constraints on participant movement, thus restricting the type of behaviour that can be investigated. An emerging alternative is wearable MEG with OPMs, enabling free movement while maintaining the millimetre and millisecond precision afforded by cryogenic MEG (Boto et al., 2018; Tierney et al., 2019). This allows recording meaningful neural signals from entire brain during walking (Mellor et al., 2023; Seymour et al., 2021), dancing (O’Neill et al., 2025), and even realistic dyadic interaction (Holmes et al., 2023).

However, the full potential of this wearable technology is not achieved with stationary visual input on screens. Because many realistic environments cannot be implemented due to the relatively small size of the required magnetically shielded room (MSR), immersive VR could provide a fully controllable method for simulating arbitrary task environments. The potential of VR has already been demonstrated for other recording modalities and model species across neuroscience domains (Alexander et al., 2024; Huang et al., 2020; Judák et al., 2025; Levy et al., 2025; Parsons, 2015; Thurley, 2022). This includes scenarios that would be difficult to implement in reality for ethical or practical reasons (Brookes et al., 2024; Sporrer et al., 2023). Immersive VR delivers visual and auditory information that depends in real-time on the movements of the participant (Judák et al., 2025; Steuer et al., 1995). This sensory input is typically transmitted to the participant via a HMD. However, consumer-grade HMDs do not currently integrate with OPM, as their electromagnetic emissions are sufficiently strong to drive OPM sensors outside their dynamic range, and to mask signals from the brain. Foundational work in this field demonstrated the ability to use OPMs in conjunction with VR but, despite removing extraneous metallic components from such a commercial HMD, the sensors could only detect signals from the most posterior parts of the occipital cortex (Roberts et al., 2019).

To overcome this gap, we designed a novel research platform that enables VR, including real-time tracking of head and hand movements through space, during OPM-based MEG. At the heart of this platform is a low-noise HMD, which is integrated with 6-degrees-of-freedom optical motion capture (MoCap) and a game engine. After designing this platform, our key goal was to validate its compatibility with the OPM-based MEG system under conditions useful and applicable to cognitive neuroscience research. On this basis, we sought to (i) assess AC and DC noise levels in the absence of a human participant, thereby quantifying potential magnetic interference introduced by the head-mounted-display (HMD); and (ii) verify the presence of well-established task-based neural responses, including those predicted to be most susceptible to HMD-induced noise due to their anatomical proximity to the HMD electronics (e.g., prefrontal cortex).

For validation experiments involving neural responses, we selected tasks that elicit measurable response robustly in individual participants. While individual differences exist even in these tasks, we were primarily interested in a proof-of-principle that the operation of the HMD does not preclude recording neural responses across various brain areas, for which a small number of participants is sufficient.

## Material and Methods

### Design of a VR system

Figure 1 shows the VR system, encompassing (1) a HMD to present dynamic visual stimuli based on movement of the head and of other body parts, (2) a MoCap system to detect the position of the head and other body parts (3) software to integrate motion information with the cognitive task, and present it in the HMD. All software and hardware components used in the development of the Optically Pumped Magnetometer Virtual Reality (OPMVR) system are listed in Supplementary Tables 1 and 2.

**Figure 1.**
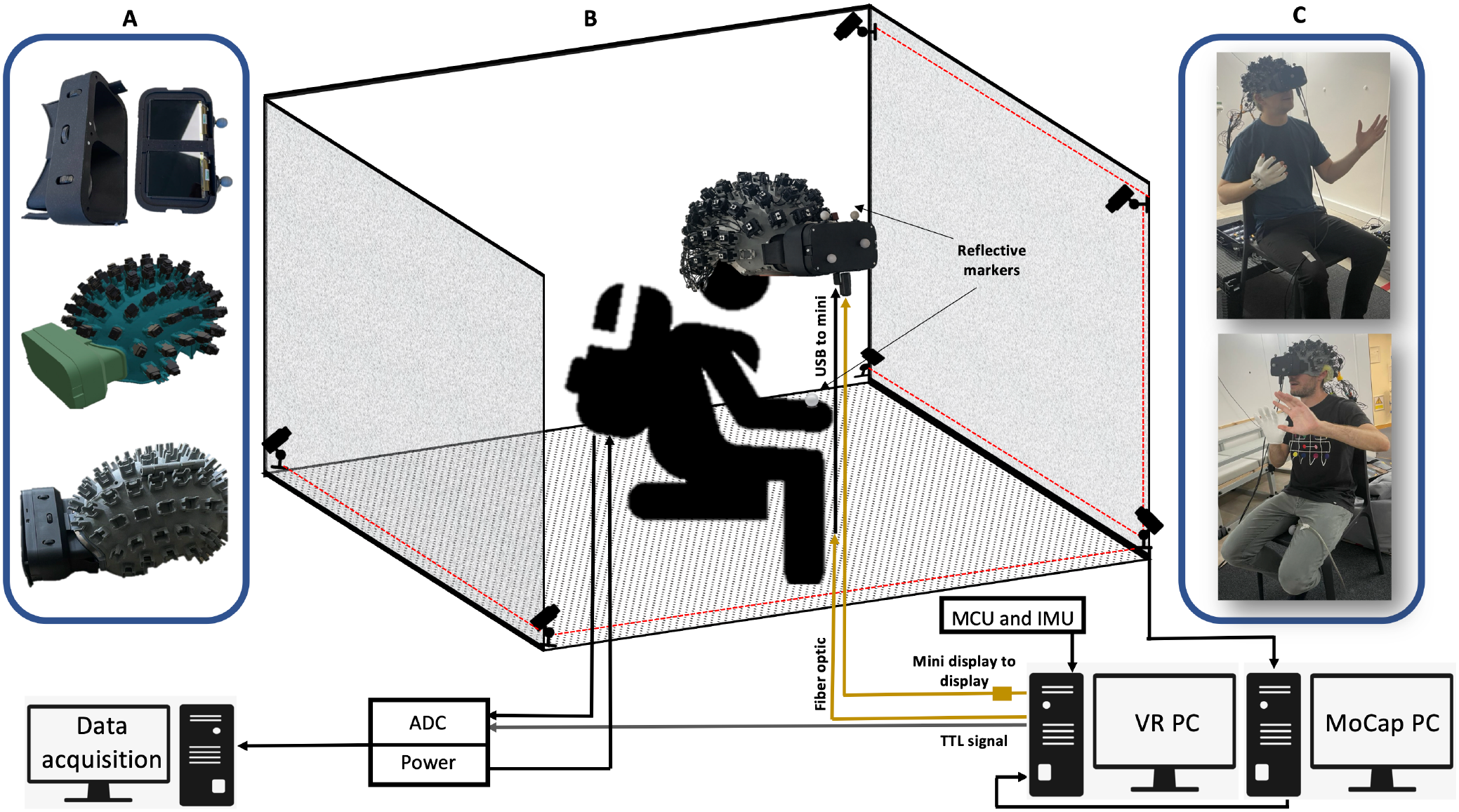
Schematic of the VR system. **(A)** Design and integration of the virtual reality headset with the OPM helmet. The displays drivers are placed behind the displays in the black plastic box (first row). The CAD model of the headset (middle row) and the 3D printed model (last row). **(B)** Schematic overview of the experimental room setup. The participant is seated inside a magnetically shielded room equipped with 6 motion capture cameras. The HMD is connected via fibre-optic USB and display cables to the VR PC. The motion capture PC receives camera recordings, computes the position of the markers on the hand and on the headset and transfers them to the VR PC. **(C)** Photographs of two participants (with their consent) wearing the integrated HMD, demonstrating the setup used for cognitive and motor experiments. NOTE: Panel C shows two of the authors demonstrating the VR setup. (See supplementary figure 1 for higher a resolution figure)

### Head mounted display

To ensure compatibility with popular VR applications, such as Unity (Unity Technologies, San Francisco, CA, USA) and Unreal Engine (Epic Games, Cary, NC, USA), we designed our HMD to integrate with SteamVR (Valve Corporation, Bellevue, WA, USA) through OpenVR API (Valve Corporation, https://github.com/ValveSoftware/openvr). We achieved this by following the 3 Degrees of Freedom (3-DoF) open-source HMD design developed by Relativty (https://github.com/relativty/Relativty).

#### Display

The primary components of a HMD are two display panels which independently present visual input to each eye, enabling presentation of a three-dimensional scene via binocular vision. To maximise presence in virtual reality, we sought to use a high-quality display, as defined by resolution and field of view. In general, two types of display are used for this purpose: liquid crystal display (LCD) panels, and organic light-emitting diode (OLED) panels. OLED technology relies on self-emissive pixels (i.e., each pixel is a light source, driven independently). In contrast, LCD pixels are not emissive, and rely on a backlight, which is typically provided by an always-on LED. For both display types, the current (and thus, potential electromagnetic interference) depends on the content being displayed, which in turn depends on the participant’s head movements and is thus hard to predict. Although we did not conduct a systematic comparison, we reasoned that this type of interference might be inherently stronger in OLED panels due to their design. This is supported by earlier reports (Roberts et al., 2019) of stimulus-related MEG signals arising from an OLED display panel. Therefore, we opted for a LCD panel.

In line with consumer-grade HMDs, we used dual 1440 x 1440 resolution LCD panels (Wisecoco, LCD Display, 120 Hz, 2.9 inch, MIPI Display 1440 x 1440, DP Controller Driver Board LS029B3SX02, DP to MIPI, Wisecoco, China), driven at 120 Hz. This frame rate enables a motion-to-photon latency of around 8 ms, considered adequate for comfortable VR presentation during movement (LaValle et al., 2014; Steuer et al., 1995).

#### Lenses

Following typical HMD design, we placed lenses (G11 VR Shinecon, Shinecon, China) in front of the displays to manipulate focal distance. These lenses can be adjusted laterally (left/right) and longitudinally (forward/backward) relative to the user’s head, such that inter-ocular distance could be adjusted, promoting user comfort (Cheng et al., 2021), and enabling the device to be used by multiple participants. Each lens had a diameter of 40 mm and a focal length of 55 mm.

#### Housing (headset)

A 3D-printed housing unit (Chalk Studios, London, UK) supported the display, display driver board, and lenses. Two adjustment dials were added to control the lens-to-lens and lens-to-eye distance to modify the interpupillary distance and lens-to-eye spacing, ensuring proper optical alignment for different users and comfortable viewing. The HMD was designed to be usable for different participants, by mounting it onto their participant-specific OPM sensor helmets (Figure 1A and supplementary figure 1) with two clips. Clips were designed for easy operation and can be mounted/dismounted by the participants themselves. Due to the rigid nature of both the helmet and the HMD housing, OPM sensor positions remain fixed in relation to sources of interest (i.e. the brain) and proximate sources of noise (i.e. HMD electronics).

#### Cabling

Displays were connected to their driver board via two 5 cm flexible flat ribbon cables. The mini DisplayPort on the driver board was then connected to the VR computer via a 30 m mini DisplayPort fibre-optic cable and a mini DisplayPort to DisplayPort adapter (Lindy, Thornaby, UK). The micro-USB port on the driver board was connected to the VR computer via a 3 m micro-USB to USB cable and a 10 m USB 3 fibre optic extension. Cable length was partly determined by market availability.

#### Microcontroller and IMU

OpenVR/SteamVR expect input from an Inertial Measurement Unit (IMU). To minimise electromagnetic interference, we sought to replace the IMU functionality by MoCap, but the IMU input was still required to operate OpenVR. Hence, we located a stationary IMU (MPU-6050, TDK InvenSense, San Jose CA, USA) and a ATmega32U4-based microcontroller (DIYmore Pro Micro 16 MHz, Mongkok Kln, Hong Kong, China) outside the MSR. The IMU was connected to the microcontroller through VCC, GND, SDA, and SCL pins. The microcontroller continuously streams IMU signals to emulate a valid VR tracking device, thereby satisfying OpenVR requirements while preventing electromagnetic interference inside the MSR. Programming of the microcontroller and its communication with the IMU was adapted from the Relativty open-source project (see more details in https://github.com/relativty/Relativty).

#### Software and driver

For software integration, an Arduino IDE (Arduino, Turin, Italy, https://www.arduino.cc/en/software) was programmed using the FastIMU library (https://github.com/LiquidCGS/FastIMU) for calibrating and streaming IMU data to SteamVR (Valve, Bellevue WA, USA). A custom SteamVR driver enabled real-time positional alignment and display rendering from MoCap data (see below). Parameters such as display alignment, lens distortion correction, and interpupillary distance were configured to align with SteamVR’s display coordinate system, ensuring the VR environment was accurately represented. SteamVR and the corresponding driver were installed on the VR PC.

### Motion capture system

To minimise electromagnetic interference, we did not use the IMU for motion tracking and instead relied on an optical MoCap system. An array of six OptiTrack Flex13 (NaturalPoint Inc., Corvallis OR, USA) MoCap cameras were used. These cameras were placed in corners of the room, with two in each (one high up, one low down), to allow for complete coverage of the tracking area. Four retro-reflective markers were attached to the HMD and another four to a right-hand glove in fixed positions such that either array formed a unique rigid body. These were tracked passively using the MoCap cameras at 120 Hz throughout the experiment. By measuring the joint translation of markers on these rigid bodies, the MoCap system calculates their position and orientation using Motive (NaturalPoint Inc., Corvallis OR, USA), run on a separate MoCap PC. Position and translation of these rigid bodies were streamed in real time to Unity software.

### VR control

To design and run the VR, we used Unity Engine with SteamVR, together with the open-source Unity Experiment Framework version 1 (https://github.com/immersivecognition/unity-experiment-framework) (Brookes et al., 2020) and the VRthreat Toolkit for Unity (Brookes et al., 2024). Experimental stimuli were rendered and controlled by a computer with dedicated graphics (VR PC) with Windows 10, a NVIDIA GeForce RTX 2080 Ti GPU, a i7 9700k CPU and 64 GB RAM. VR PC and MoCap PC communicated through a network connection using an ethernet cable and the OptiTrack plugin (https://docs.optitrack.com/plugins/optitrack-unity-plugin) for Unity software. To synchronise VR and OPM data acquisition, the VR PC sent TTL markers via its parallel port.

### OPM system

The experiments took place in a MSR (Magnetic Shields, Ltd., Staplehurst, UK), with internal dimensions of 3 x 4 x 2.2 m. Before the experiment began, the room was degaussed to minimise residual magnetic fields. OPM sensors (QuSpin Inc., Louisville CO, USA) were secured in sockets within a rigid sensor helmet, custom-built based on each participant’s T1-weighted structural 3T MRI (Meyer et al., 2017). This design ensures precise co-registration, optimises signal quality for various head sizes, and minimises sensor, cable and HMD movement relative to the head (Figure 1A).

The Neuro-1 acquisition system (QuSpin Inc., Louisville CO, USA) was used, incorporating 64 tri-axial sensors operating in closed-loop mode. The sampling frequency was 1500 Hz for participant 1 and 375 Hz for participant 2, following a software update that changed the default acquisition settings. The OPM sensors (QZFM Gen3) have an intrinsic bandwidth of 0-135 Hz, determined by the properties of the vapor cell. Additionally, the system included a 6^th^ order digital low-pass Chebyshev (type 1) IIR filter at 150 Hz, implemented by the manufacturer.

### Experimental validation of the virtual reality system

We used two strategies to validate our VR system. First, we assessed DC and AC noise contributions of the HMD system in the absence of a human participant, in order to evaluate the potential for electromagnetic interference with the OPM sensors. Second, we used an experiment-based calibration approach (Bach et al., 2020), where the validation criterion is to reproduce well-known observations with a new measurement system. We used experiments that are established to elicit specific neural responses, and evaluated whether these were observed when using our VR system (Figure 2). Thus, we sought to demonstrate that any electromagnetic interference is small enough to render the VR system practically useful.

**Figure 2.**
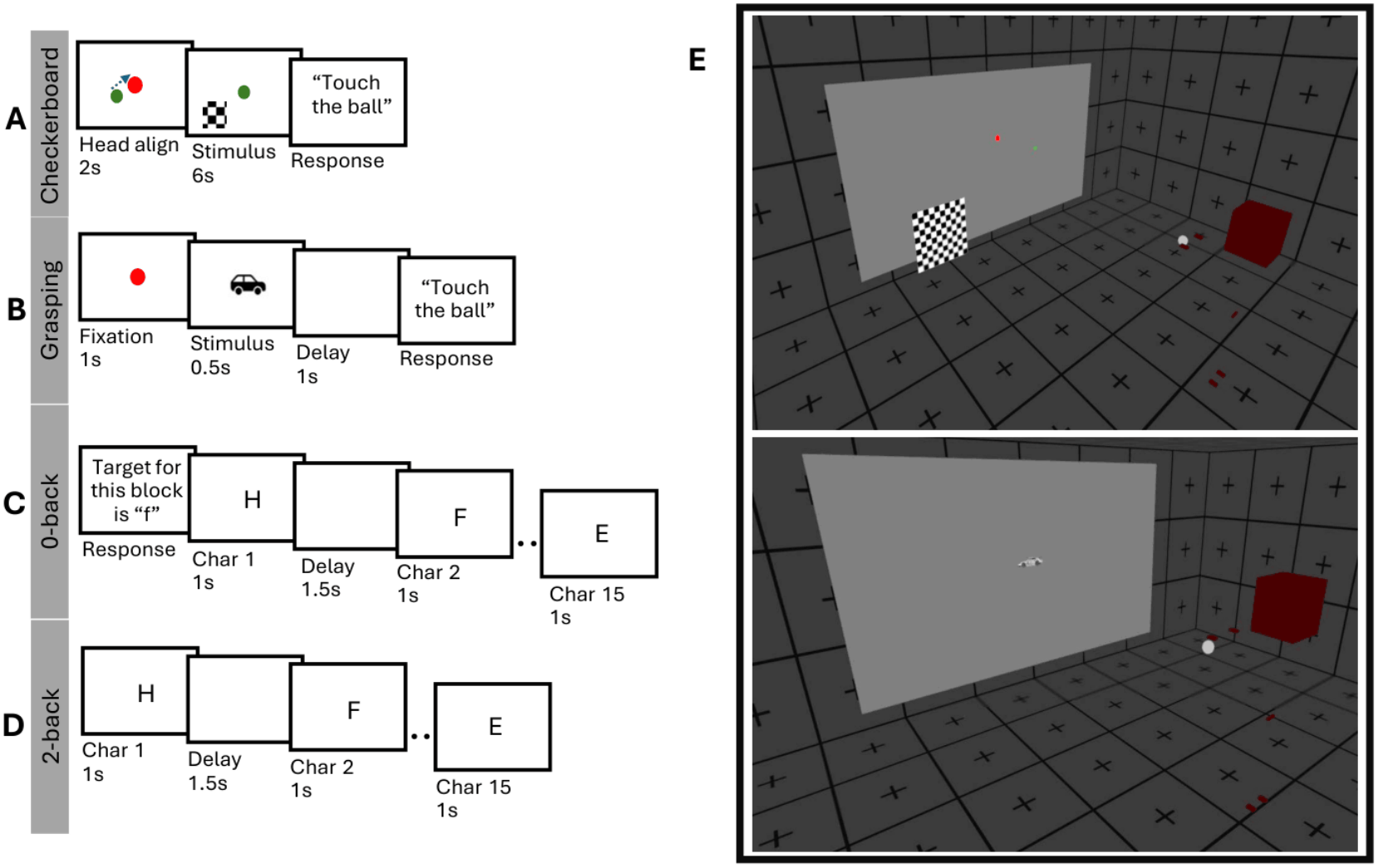
Schematic overview of cognitive and sensorimotor tasks performed with participants during MEG recordings in virtual reality environment. **(A)** Checkerboard task: Participants aligned their head position using a fixation dot, then initiated the trial by touching a response ball. A dynamic checkerboard stimulus (4 Hz, 6 s duration) appeared 8.5° off-centre to evoke visual responses. **(B)** Grasping task: Participants fixated on a red dot and grasped the response ball to initiate the trial. After a 1 s delay, a face or car image was presented for 0.5 s, followed by a 1 s delay and a prompt to repeat the grasping action, targeting motor activity. **(C)** 0-back task: Participants viewed a sequence of letters (1 s presentation, 1.5 s delay). They responded by touching the ball when a letter matched a predefined target shown at the block’s start. **(D)** 2-back task: Similar to the 0-back, but participants responded only if the current letter matched the one presented two trials before, engaging working memory processes. **(E)** screenshots of the virtual environment for the checkerboard (up) and grasping experiment.

#### Participants

Our goal was to demonstrate the possibility of recording neural signals from the entire brain in the presence of our HMD, rather than address the consistency or variability of the neural signals across participants. We recruited two participants (33 and 30 years, male), who underwent two recording sessions on different days. The experiments were conducted in accordance with the Declaration of Helsinki, and were approved by UCL Research Ethics Committee (3090/005).

### Noise measurement

#### Experiment 1: DC noise

To quantify potential interference from the HMD, we measured the magnetic flux density generated by metallic components (e.g. screen driver board) at different distances from the HMD, using a single-axis fluxgate magnetometer (FLUXMASTER, Stefan Mayer Instruments, Dinslaken, Germany; resolution 0.1 nT, range ±2 to ±200 µT). The HMD was turned off during these measurements. We first recorded a measurement of the baseline magnetic field with the HMD positioned 1 meter away, while the front face of the HDM was pointing toward the sensor (noting that beyond this distance no further change in magnetic flux was observed), followed by repeated measurements against this baseline with the HMD introduced at closer distances (50 cm, 25 cm, 12.5 cm, 6.25 cm, 3.125 cm). As the fluxgate magnetometer was single-axis, measurements were repeated with the sensor in three orthogonal orientations, while keeping its position fixed. We report the magnitude of the magnetic flux density by taking the Euclidean norm of those three orthogonal measurements at each distance, and subtracting the baseline (magnetic flux when the headset is at the 1 m distance) magnetic flux density magnitude for display purposes.

#### Experiment 2: AC noise

To test whether a well-controlled magnetic dipole-like signal could be reliably detected and localised in the presence of AC noise from the HMD, we placed a coil connected to a signal generator inside the OPM helmet, which hosted 64 sensors. The coil emitted a 30 Hz sinusoidal oscillatory magnetic field (a magnetic dipole source), while OPM signals were recorded. The phantom signal was then localised by fitting an equivalent current dipole to the averaged field (using FieldTrip dipole fitting with infinite homogeneous head model). In two experimental conditions, HMD which was attached to the helmet, either displaying a windows desktop screen or turned off (1 minute off, 1 minute on, one minute off). Both helmet and the HMD were placed in the centre of the room.

### Basic stimulus evoked response experiments

#### Experiment 3: Eyes open vs. eyes closed

Similar to previous HMD evaluation studies (Hertweck et al., 2019; Roberts et al., 2019), participants were instructed to keep their eyes open for one minute, and then closed for another minute, while the HMD was displaying a Windows desktop screen. Instructions were delivered via a microphone from outside the magnetically shielded room and played through a loudspeaker, also outside the room. In line with previous literature (Barry C De Blasio, 2017; Hohaia et al., 2022; Roberts et al., 2019), we expected higher alpha (8-12 Hz) activity in the occipital areas overall, as well as increased power when eyes were closed (i.e., the classic Berger effect (Berger, 1929)).

To localise the source of this effect, the continuous recording was divided into consecutive non-overlapping epochs of 2 s, with epochs from the first minute labelled as the eyes-open condition and those from the second minute as the eyes-closed condition. Because no explicit trigger marked the eyes-open–to–eyes-closed transition, the two epochs spanning the period immediately around the verbal instruction were excluded to avoid contamination from the transition. Source localisation was then performed in the alpha band (8–12 Hz) using an LCMV beamformer contrasting eyes-closed against eyes-open power.

#### Experiment 4: Flickering checkerboard

Similar to previous HMD evaluation studies (Hertweck et al., 2019; Roberts et al., 2019), we assessed visual cortex entrainment to a flickering checkerboard. Participants were instructed to align their head position so that a green dot—controlled by their head movement—overlapped with a red dot displayed in the centre of the screen in front of them in the virtual environment. They were then asked to maintain visual fixation on the red dot throughout the stimulus presentation. They were asked to touch the white virtual response ball which placed in front of them at less than arm distance with the right hand whenever they fixated on the red dot and were ready for the next trial. In each trial, after a 2 s delay, a 4 Hz alternating checkerboard pattern was presented for 6 s, 8.5° off-centre from the fixation point at a size corresponding to 9° of visual angle. Immediately after the stimulus, the instructions to touch the response ball appeared. The experiment consisted of two blocks of 60 trials (Figure 2A). Extrapolating from well-known entrainment effects (Herrmann, 2001; Jiang et al., 2007), we expected 4-12 Hz activity (including the base frequency and two harmonics) in contralateral visual cortex, and applied small-volume correction within an a priori bilateral calcarine (primary visual) cortex region of interest, as defined by the Neuromorphometrics atlas in SPM (labels 108 and 109).

#### Experiment 5: Grasping

We assessed motor cortex activity related to a motor task that was embedded in an incidental cognitive task to maintain attention. As in experiment 4, participants were asked to fixate on a red dot in the centre of the screen and then start the trial by grasping the response ball. After grasping, with a delay of 1 s, a face or car image were shown on the screen for 0.5 s. Following the disappearance of the image, a text appeared with 1 s delay and asked the participant to grasp the ball again to start the next trial. The experiment consisted of two blocks of 101 trials (Figure 2B). Based on previous reports (Jurkiewicz et al., 2006), we expected beta-band (13-30 Hz) activity localized to motor cortex, along with a power decrease in this frequency band associated with motor preparation and execution. We expected motor-related activity to localize to the left precentral and postcentral gyri (Jurkiewicz et al., 2006) and therefore applied small-volume correction using Neuromorphometrics-defined (a built-in atlas in the SPM package) regions of interest in SPM (labels 183 and 177).

### Cognitive experiments

#### Experiment C: N-back test

In this working memory experiment, participants completed an N-back task. They were asked to fixate on a red dot on the screen and started each trial by grasping a response ball. After a 1 s delay, letters were sequentially presented at the centre of the screen, each lasting 1 s with a 1.5 s gap between presentations. Depending on the N-back condition (0-back, or 2-back), participants were instructed to touch the ball when the current letter matched either a previously presented letter (2 steps prior) or a target letter shown at the start of the block (0-back). After 15 trials (i.e., letters), a rest period followed, and participants initiated the next block of 15 trials by touching the ball. The experiment contained 20 blocks of 15 trials (300 trials in total), and there was an obligatory break after half of the blocks. Task instructions and feedback were provided through on-screen prompts (Figure 2C and 2D). Since frontal-midline theta-band (4–8 Hz) activity during working memory localises to medial frontal cortex (Brookes et al., 2011; Rhodes et al., 2023), we restricted our analysis to this band and, a priori, to a medial frontal region of interest. Small-volume correction was applied within the medial superior frontal gyrus as defined by the Neuromorphometrics atlas in SPM (labels 152 and 153).

### General procedure

For the DC and AC noise experiments, no human participants were involved. In the DC experiment, two researchers inside the MSR adjusted the distance of the HMD from a single-axis fluxgate magnetometer and measured the resulting magnetic fields. In the AC experiment, the HMD together with an OPM sensor helmet with all sensors mounted on it, and a coil placed inside the helmet were set up inside the MSR. No person was present in the MSR during the recording.

For experiments involving neural responses, participants were first guided to sit on a chair inside the MSR to mount the OPM sensor helmet. After putting the MoCap glove on their right hand, they attached the HMD. They were asked to repeatedly remove and reattach the HMD to familiarise themselves with the procedure. Participants were also given an emergency buzzer (Figure 1C).

Before each individual experiment started, participants were located in an empty virtual room with grey walls. The virtual wall in front of them carried a virtual screen displaying task instructions and visual stimuli. In experiments that required participants’ response, there was a white virtual response ball, which participants could touch or squeeze with the gloved hand.

Once the setup was complete, all researchers exited the room, and the MSR was sealed. From this point onward, all communication with participants occurred via a two-way intercom system. In experiment 3, instructions were delivered verbally through the intercom. For all other experiments, instructions were displayed on the virtual screen inside the VR environment.

### Analysis

Analysis was performed using SPM25 (Tierney et al., 2025) and FieldTrip (Oostenveld et al., 2011) in MATLAB (R2021b).

### Pre-processing

MEG data were imported into SPM, and downsampled to 375 Hz after applying an anti-aliasing low-pass filter. Power spectral analysis was used to visually detect faulty channels, identified by substantial deviations from the median power or the manufacturer’s noise floor. For experiment 2 (AC noise) 46 channels were excluded. We use the terminology ‘channel’ to describe each of the sensitive axes of the triaxial OPM sensor. In the other experiments for participant 1, 15 channels were excluded in session 1 and 26 in session 2; for participant 2, 14 and 17 channels were excluded, respectively.

Data were additionally high pass filtered at 2 Hz and and line noise was removed using Zapline-plus (Klug C Kloosterman, 2022), which attenuates the 50 Hz mains and its harmonics via spectral–spatial filtering. Adaptive multipole models were applied to reduce environmental noise across a broad frequency range (Tierney et al., 2024). For analysis of specific frequency bands, data were further filtered to the specific band of interest.

Where appropriate, the data were segmented into individual trials based on event markers. The specific time windows analysed are reported in the results section.

### Time-frequency analysis

Time-frequency analysis was performed for Experiment 3 to quantify changes in alpha-band power (8–12 Hz), given its well-established modulation by eye state. Continuous MEG data were analysed using Morlet wavelet decomposition implemented in SPM over a two-minute recording period (one minute eyes open followed by one minute eyes closed), yielding time-resolved estimates of alpha-band power. Power values were normalised to the average power during the initial eyes-open period and expressed in decibel (dB) units. Topographic maps were generated to illustrate the spatial distribution of alpha power across the sensor array, and representative sensor time courses were selected based on the sensors exhibiting the highest average alpha power across the entire recording.

### Source localisation

Source localisation analysis was conducted using the DAiSS toolbox in SPM. Linearly constrained minimum variance (LCMV) beamformer weights were computed within time windows and frequency ranges specific to each analysis, with these parameters detailed separately in the results section.

The source space was defined as a 3D grid spanning the entire brain volume, constrained by the inner skull, with a spatial resolution of 5 mm. Sensor locations and orientations were inherently registered to MRI space, as each participant’s sensor helmet was created based on their individual MRI scan (Meyer et al., 2017). To ensure consistency and facilitate interpretation, data were transformed into canonical Montreal Neurological Institute (MNI) space within SPM. This transformation was achieved via an affine alignment between fiducial markers in native space and MNI coordinates. Source orientation was optimised to maximise signal power.

For each trial and condition, a volumetric power image was generated and smoothed with an 15 mm isotropic FWHM Gaussian filter (Gross et al., 2013).

### Statistical analysis

Statistical inference used voxel-wise permutation testing (Nichols C Holmes, 2002). Condition labels were permuted across trials over 10,000 permutations; on each permutation the maximum F-statistic across the search volume was retained to form the null distribution of the maximum statistic, giving FWE control at the voxel level (p < 0.05). All analyses used small-volume correction: the search volume was restricted to an a priori region of interest defined from Neuromorphometrics atlas labels (separately for each task). Permutations shuffled trial labels within each participant. For display, voxels surviving FWE correction are shown as the signed t-statistic, with the sign indicating the direction of the effect. Figures thresholded at half-maximum are provided in the Supplementary Material.

### Dipole fitting

For experiment 2 with a 30 Hz phantom source, we used magnetic dipole fitting for source localisation. The signal underwent the same preprocessing described above (low-pass filtering at 90 Hz, high-pass filtering at 2 Hz, line-noise removal at 50 Hz using Zapline, and AMM) and was then band-pass filtered between 25 and 35 Hz and segmented into 50 ms epochs (time-locked to the phantom signal using a reference recording of the signal generator output driving the coil). We then averaged over epochs and applied source localization to estimate the origin of the 30 Hz signal within the helmet. We identified the peak of the signal by finding the time point with the maximum absolute magnetic field amplitude across all magnetometers. A short time window around this peak (two consecutive time samples, peak time and peak time +1, or peak time and peak time -1 if the peak occurred at the final time point) was selected for dipole fitting. We then performed a grid-based dipole fitting using the Fieldtrip function *ft_dipolefitting,* assuming a magnetic dipole in an infinite homogenous medium, and with grid search enabled to identify the best-fitting dipole within the head model during the specified time window. The estimated source locations were compared between conditions to determine whether the HMD introduced localisation bias.

### Code and data availability

All code and data will be made publicly available on OSF (https://osf.io/qpx37).

## Results

### Experiment 1: HMD generates minimal magnetic flux

Figure 3A shows the magnitude of magnetic flux density at distances between 3.125 cm and 50 cm, encompassing those in our HMD design, where OPM sensors are located more than 5 cm away from the HMD. To put these into context, we compare them from a commercially available HMD (Pimax 5K Plus, Pimax Innovation Inc., Shanghai, China) and the same HMD with non-essential metal components (e.g. screws) removed. As expected, the impact of the proposed HMD on the magnetic field is approximately one order of magnitude smaller than for the de-metalled HMD, and two orders of magnitude smaller than for a commercial HMD. Crucially, we found that the field was smaller than 18 nT when the HMD was at 31.25 mm distance (see Figure 3A), which is within the dynamic range of commercially available OPM sensors when operated on closed-loop mode (Yan et al., 2022).

**Figure 3.**
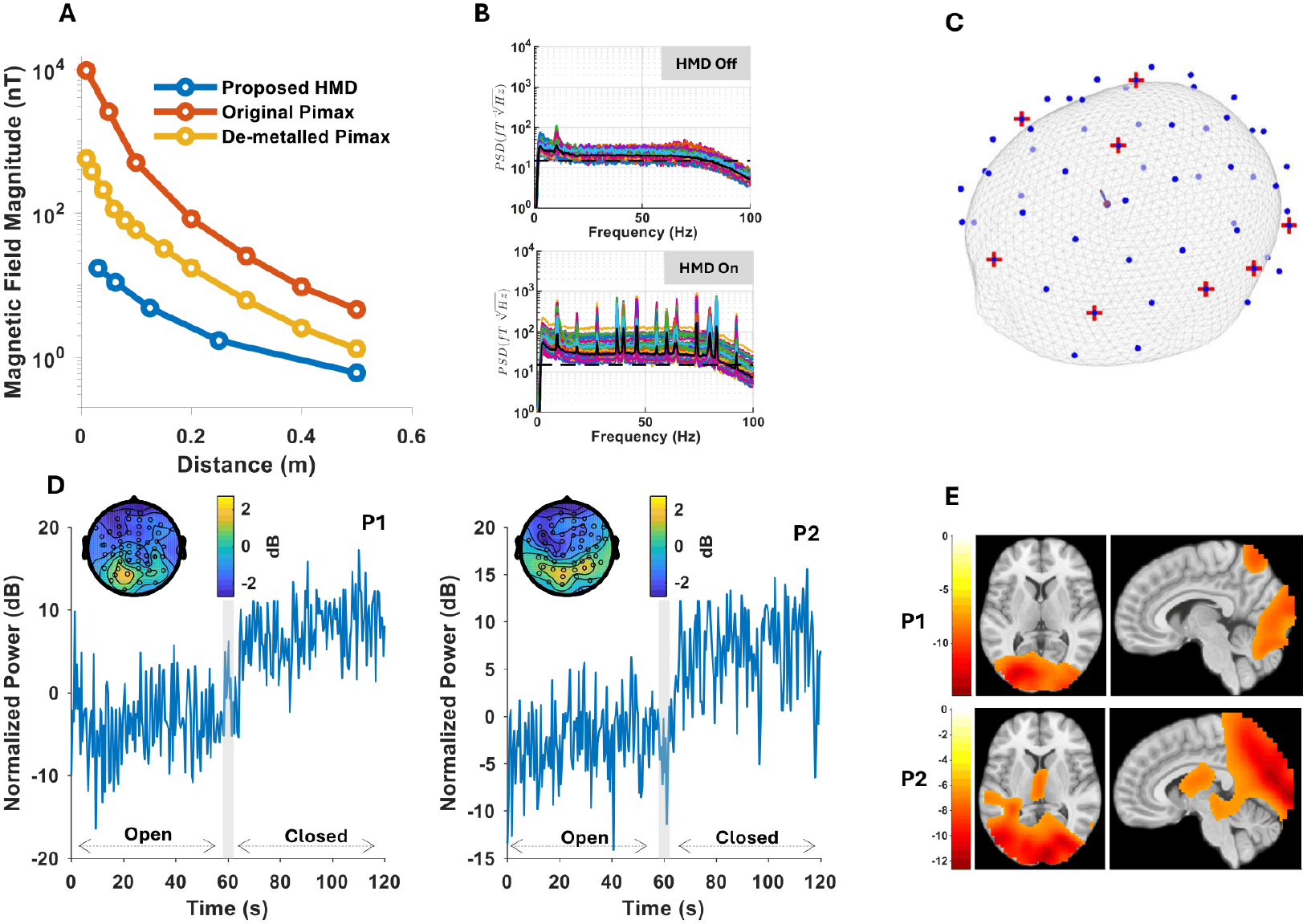
Characterisation of magnetic interference from the VR headset and the power of different frequency bands over time in eye-opened and eye-closed conditions. **(A)** DC magnetic flux magnitude measured as the Euclidean norm of three orthogonal measurements at each distance, and subtracting the baseline magnetic flux density, shown for the proposed MEG-compatible HMD (blue), a commercially available Pimax headset (red), and a de-metalled version of the Pimax (orange) across multiple distances. **(B)** Power spectral density (PSD) of the preprocessed OPM data, with the HMD off (top) and on (bottom). Each coloured trace is a single OPM channel; the bold black line is the channel median and the dashed horizontal line marks the 15 fT/√Hz reference. **(C)** Source localization plots show the estimated origin of the 30Hz signal during headset-off (blue arrow) and headset-on (red arrow), which differed by 0.14 mm between conditions. Sensors with out-of-range power spectra (i.e., bad channels, see pre-processing for details) are marked in red crosses on the schematic head surface and were distributed across the entire helmet, not concentrated in regions close to the HMD (see supplementary figure 1 for the position of bad sensors in other experiments). **(D)** MEG alpha-band power from the sensor exhibiting the highest average alpha power across the experiment (subsampled by a factor of 10 for clearer visualization). Gray bar shows the time range when participants are instructed to close their eyes. The insets illustrate the topographic distribution of alpha power averaged over all time points. As shown in the map, the sensor with the maximum average alpha power is located over the occipital region (yellow region) for both participants. During the experiment, participants were verbally instructed via microphone to keep their eyes open for one minute, followed by closing their eyes for another minute. **(E)** Source localisation of the alpha-band (8–12 Hz) eyes-closed versus eyes-open contrast for both participants, confirming that the increase in alpha power during eye closure originates in occipital cortex. Maps show the signed t-statistic within voxels surviving whole-brain family-wise-error correction (p < 0.05), assessed by a two-tailed permutation F-test (10,000 permutations); for visualisation, only significant voxels exceeding half the peak value are displayed. Colour bars indicate t-statistic values.

### Experiment 2: Phantom signal localisation error is minimal

To characterise potential HMD interference, we first recorded the OPM signal with the HMD turned on and off. After preprocessing (2 Hz high-pass, 90 Hz low-pass, line-noise removal and AMM), switching the HMD on raised the broadband noise floor by ∼3.1 dB (from 20.0 to 28.5 fT/√Hz, taken as the median of the channel-median amplitude spectral density across 5–90 Hz) and introduced a series of discrete narrowband peaks, most prominent at around 40, 46, 60, 74 and 83 Hz, up to ∼16 dB above the local noise floor, relative to the headset-off condition (Figure 3B; see Supplementary Figure 3 for the spectra before AMM). These narrowband peaks likely reflect electromagnetic emissions from active electronic components of the headset, including display drivers and data transmission processes, whose operating frequencies and harmonics introduce structured spectral artefacts.

To test whether this interference affects source reconstruction, we then placed a 30 Hz magnetic phantom (an internal coil) inside the OPM helmet and localised it with the HMD turned on and off. Despite the presence of these spectral features, the source localisations of the magnetic phantom differed by 0.14 mm between the two conditions. Thus, AC noise generated by the HMD had no appreciable impact on source localisation at this frequency. Importantly, the out-of-range sensors (i.e., bad channels, see pre-processing for the bad channel selection process) were not clustered near the HMD, suggesting that their signal quality issues were not caused by the HMD itself (Figure 3C, see supplementary figure 2 for the position of the out-of-range sensors in other experiments).

### Experiment 3: Increased occipital alpha activity during eye closure

Figure 3D insets show the topographic map of alpha (8-12 Hz) amplitude in the radial OPM channels, which is maximal in the occipital area as expected. The extracted alpha power time series from the sensor with the strongest average alpha activity shows a clear increase in the amplitude of power alpha (8-12 Hz) band for both participants when they are instructed to close their eyes. Figure 3E shows the source localisation of the eyes-closed versus eyes-open contrast, computed across the whole brain and corrected for multiple comparisons. As expected, the increase in alpha power localised to occipital cortex. For visualisation, only significant voxels exceeding half of the peak value are displayed.

### Experiment 4: Visual cortex responses induced by dynamic checkerboard

MEG signal was recorded while a lateralised 4 Hz (8 reversals per second) dynamic checkerboard pattern was displayed on either side of the fixation point (see Figure 2A). Participants gave responses with the right hand. Source localisation revealed contralateral visual cortex responses in for participant 2 in response to both left and right visual field stimulation and for participant 1 (possibly due to lack of fixation) only in response to left visual field stimulation (Figure 4A; two-tailed permutation F-test, FWE-corrected within an a priori calcarine region of interest (bilateral calcarine cortex, Neuromorphometrics labels 108/109); see Supplementary Figure 4 for the whole-brain signed-t maps of each contrast direction, thresholded at half the peak value for illustration).

**Figure 4.**
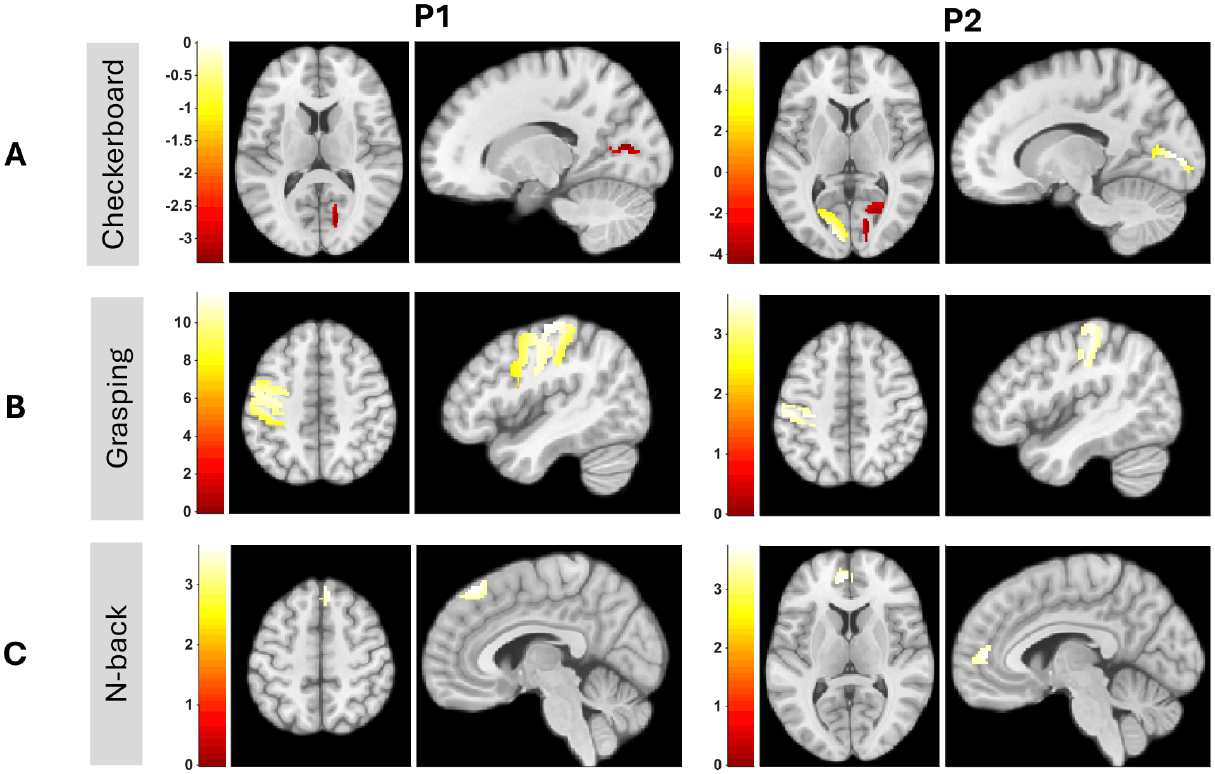
Visual, motor cortex and frontal responses across three tasks in two participants (P1, P2). Each panel shows the signed t-statistic within voxels surviving family-wise-error (FWE) correction (p < 0.05), assessed by a two-tailed permutation F-test (10,000 permutations) with small-volume correction in an a priori region of interest. Colour bars indicate t-statistic values; the sign indicates the direction of the contrast. **(A)** Checkerboard: contralateral visual cortex responses to left-vs. right-field stimulation in the 4–12 Hz band, corrected within a bilateral calcarine region of interest (Neuromorphometrics labels 108/109). **(B)** Grasping: beta-band (13–30 Hz) responses during grasp execution vs. a matched still period (1-s intervals), corrected within the left precentral and postcentral gyri (Neuromorphometrics labels 183/177). **(C)** N-back: theta-band (4–8 Hz) responses to letters in the 2-back vs. 0-back task, corrected within the bilateral medial superior frontal gyrus (Neuromorphometrics labels 152/153).

### Experiment 5: Beta-band motor cortex responses induced by grasping

Participants were instructed to grasp a ball with their right hand whenever prompted. We compared beta power (13–30 Hz) in the grasp period (1 s interval centred on grasp onset, i.e., when the hand collides with the response ball) against a matched "still" period (1 s interval centred around stimulus onset, when participants remained motionless). As expected from previous reports (Jurkiewicz et al., 2006), we observed beta-band activity in the contralateral (i.e., left) precentral and postcentral gyri associated with grasp execution (two-tailed permutation F-test, FWE-corrected within an a priori region of interest comprising the left precentral and postcentral gyri (Neuromorphometrics labels 183 and 177); Figure 4B; see Supplementary Figure 4 for the whole-brain signed-t maps of each contrast direction, thresholded at half the peak value for illustration).

### Experiment 6: Theta-band medial frontal gyrus responses induced by N-back task

Participants completed a 0-back and 2-back task (Figure 2 CD). We compared theta power (4–8 Hz) during 1 second of letter presentations between the two tasks. As expected from previous reports of frontal-midline theta power increase during working memory tasks (Jensen C Tesche, 2002; Brookes et al., 2011; Rhodes et al., 2023), we observed medial superior frontal gyrus responses in the 2-back compared with the 0-back task in both participants (two-tailed permutation F-test, FWE-corrected within an a priori region of interest comprising the bilateral superior frontal gyrus, medial segment (Neuromorphometrics labels 152 and 153); Figure 4C; see Supplementary Figure 4 for the whole-brain signed-t maps of each contrast direction, thresholded at half the peak value for illustration).

### Participants movement is much more than acceptable range in classical MEG

Participants were not instructed to sit still, as is mandatory in cryogenic MEG settings. To evaluate their actually executed movements during the experiment, we quantified the head and hand translational displacement of participants during the source localization windows for experiment 4 to 7 (Figure 6). Maximum head/hand displacement per trial was quantified as the peak translational excursion of the head/Hand tracker from its trial-onset position. Analysis of the maximum head displacement (Figure 6A, first column) per trial showed a distribution of movement that exceeded the 0.5 cm constraints typically required for traditional cryogenic MEG (Gross et al., 2013). Randomly sampled trajectories of head (Figure 6A second column) and hand positions (Figure 7B second column) also demonstrated that participants engaged in active reaching and task-based interactions without compromising the integrity of the whole-brain recordings. These results demonstrate that the proposed VR system does not require participants to remain perfectly still.

**Figure 6.**
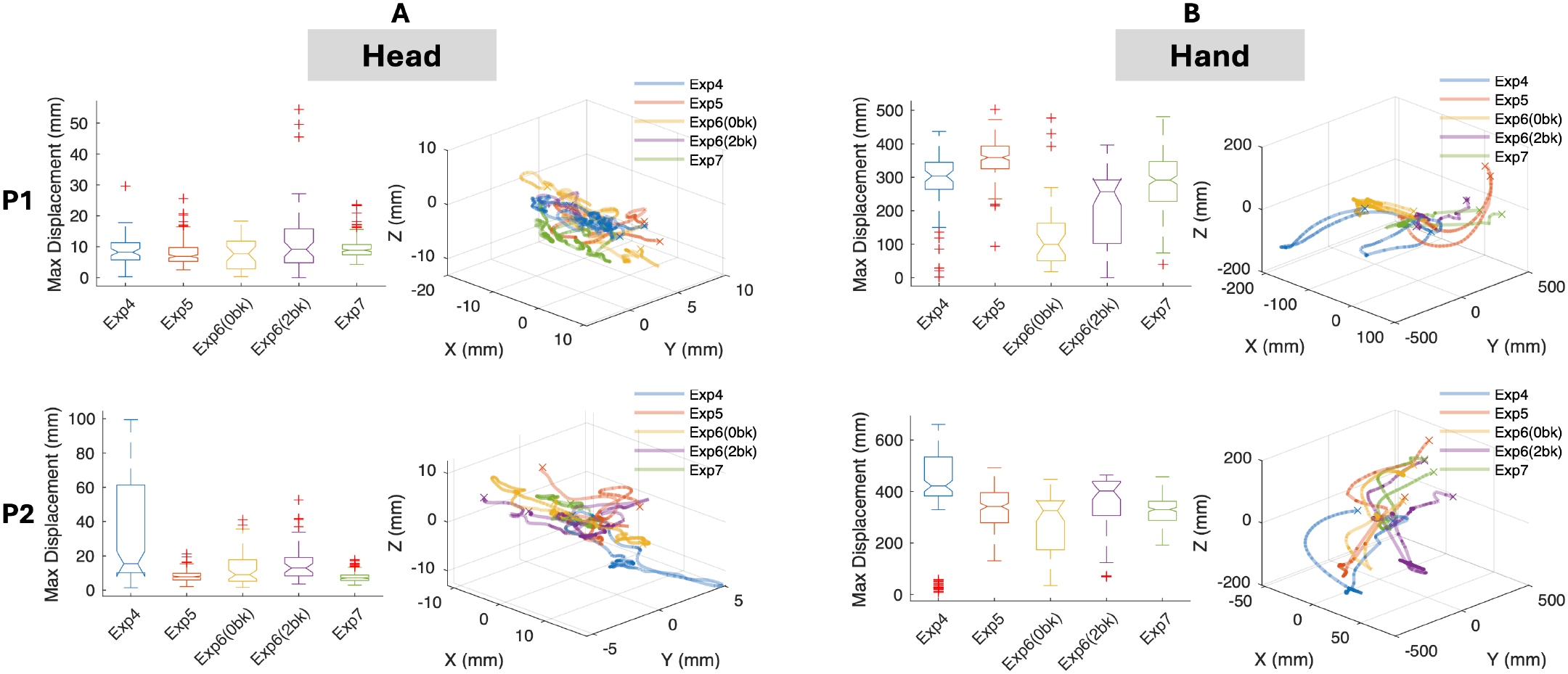
Characterization of participant movement during the time period used for source analysis. **(A)** Head movement metrics across tasks. Left column: Box plots displaying the maximum head displacement (mm) per trial for each experiment (Exp 4–7) for both participants (P1, P2). Right column: 3D trajectory plots illustrating representative head movement through space, colored by experiment. **(B)** Hand movement metrics across tasks. Left column: Box plots showing the maximum hand displacement (mm) per trial, reflecting the range of motion during manual interactions. Right column: 3D trajectories of hand movement illustrating reaching and interaction patterns within the virtual environment.

## Discussion

In this study, we present a bespoke OPM-compatible VR system which enabled neuromagnetic fields to be successfully recorded from multiple brain regions. This overcomes key limitations of using commercial VR headsets, which generate substantial magnetic interference and confine reliable OPM signal acquisition primarily to posterior brain regions. Our new system contained minimal ferromagnetic material reducing static magnetic flux density by two orders of magnitude compared to a commercial headset (figure 3A). This meant that even OPM sensors close to the eyes remained within their operational range (figure 3B). We were able to reproduce known responses in several perceptual and cognitive benchmark tasks, involving the visual, motor, prefrontal cortex and hippocampus. Thus, our results demonstrate the feasibility of novel HMD designs for OPM recordings. At the same time, the HMD offers similar latency, spatial resolution, and framerate, compared to commercial products. The HMD is based on an open-source design, adapted for OPM based MEG and is straightforward to produce. No additional components are required to navigate freely in room-scale VR.

The main limitation of our HMD design is that the display drivers are in direct vicinity to the display itself, creating residual electromagnetic interference (e.g. figure 3B). Reducing this interference would be particularly useful in order to detect neural sources of small magnitude, to improve source reconstruction across the whole brain without a priori assumptions, and to better isolate movement-induced signal artefacts (Seymour et al., 2022). For example, given the necessarily limited selection of brain areas investigated in the present report, it remains unclear whether interference generated by the HMD has a spatial structure and would potentially distort source localisation in specific areas. One reason for housing the display drivers together with the displays is that they are connected via ribbon cables that bend in one axis but cannot as easily be twisted as the other cables we used. Thus, affixing the display drivers to the participant’s body could restrict their movements. Overcoming this problem is largely a mechanical design challenge. Another reason for this design choice was to keep the display drivers in a rigidly fixed position with respect to the OPM sensors, which reduces the complexity of the interference. Designs that separate the displays from their driver boards would preferably use a large enough distance to minimise any interference, or keep their position rigidly fixed in relation to the OPM sensors.

One consideration is that our HMD remains tethered to the VR computer, which constrains fully naturalistic movement. However, this limitation does not affect current OPM recordings, which themselves rely on wired sensors. As OPM technology advances toward portable systems (Schofield et al., 2024), on-participant digitisation (Hill et al., 2025), and wireless data transmission from participant to recording PC, integrating a wireless VR interface as in various commercial HMDs will be a natural next step and does not require a re-design of the HMD. Another limitation is that our HMD does not yet integrate with commercially available sensor helmets, which could be overcome by a simple adjustment of our clip-on adapter.

Notably, while our participants were free to move their heads during recordings, we did not explicitly instruct them to do so. Whilst we do expect increased sensor noise due to movement through a non-zero field (Boto et al., 2018) (Seymour et al., 2021), we do not expect the HMD to negatively impact this (as the HMD and sensors a rigidly connected), and future work will demonstrate the feasibility of our VR system for fully immersive research.

It is worth noting that the HMD noise source (screen or display driver) can be spatially well-characterised (Tierney et al., 2024) and will be constant despite head-movement. That said, one disadvantage of having the display and brain signal in perfect registration is the potential for display information to leak into the sensors (e.g. above chance classification of task dependent on display and not brain) and experimental design must take this into account.

In summary, by establishing and validating a system for combining OPM with VR, we aim to expand the methodological toolkit available for cognitive neuroscientists, bridging the gap between high-resolution neuroimaging and naturalistic behavioural paradigms. This work lays the foundation for future research in spatial navigation (Bush et al., 2017), critical decision-making (Sporrer et al., 2023) and sensorimotor closed-loop control (Pezzulo C Cisek, 2016), exploring brain function in dynamic, interactive environments while maintaining the high data quality required for neuroscience investigations.

## Acknowledgements

This work was supported by the European Research Council (ERC) under the European Union’s Horizon 2020 research and innovation programme (DRB: ERC-2018 CoG-816564 ActionContraThreat), the Discovery Research Platform for Naturalistic Neuroimaging, funded by Wellcome (DRB, GB: 226793/Z/22/Z), and by the Ministry of Culture and Science of the State of North Rhine-Westphalia, Germany (DRB: iBehave network, and PB22-063A InVirtuo 4.0). The Hertz Chair for Artificial Intelligence and Neuroscience in the Transdisciplinary Research Area Life and Health, University of Bonn, is funded as part of the Excellence Strategy of the German federal and state governments. This article is published under a CC BY open access license. Copyright is retained by the authors.

